# Suggested role of NosZ in preventing N_2_O inhibition of dissimilatory nitrite reduction to ammonium

**DOI:** 10.1101/2023.02.16.528904

**Authors:** Sojung Yoon, Hokwan Heo, Heejoo Han, Dong-Uk Song, Lars R. Bakken, Åsa Frostegård, Sukhwan Yoon

## Abstract

Climate change and nutrient pollution are among the most urgent environmental issues. Enhancing the abundance and/or the activity of beneficial organisms is an attractive strategy to counteract these problems. Dissimilatory nitrate reduction to ammonium (DNRA), which theoretically improves nitrogen retention in soils, has been suggested as a microbial process that may be harnessed, especially since many DNRA-catalyzing organisms have been found to possess clade II *nosZ* genes and the ability to respire N_2_O. However, the selective advantages that may favor these *nosZ*-harboring DNRA-catalyzing organisms is not well understood. Here, the effect of N_2_O on Nrf-mediated DNRA was examined in a recently isolated soil bacterium, *Bacillus* sp. DNRA2, possessing both *nrfA* and *nosZ* genes. The DNRA metabolism of this bacterium was observed in the presence of C_2_H_2_, a NosZ inhibitor, with or without N_2_O, and the results were compared with C_2_H_2_-free controls. Cultures were also exposed to repeated oxic-anoxic transitions in the sustained presence of N_2_O. The NO_2_^−^-to-NH_4_^+^ reduction following oxic-to-anoxic transition was significantly delayed in NosZ-inhibited C_2_H_2_-amended cultures, and the inhibition was more pronounced with repeated oxic-anoxic transitions. The possible involvement of C_2_H_2_ was dismissed since the cultures continuously flushed with C_2_H_2_/N_2_ mixed gas after initial oxic incubation did not exhibit a similar delay in DNRA progression as that observed in the culture flushed with N_2_O-containing gas. The findings provide novel ecological and evolutionary insights into the oft-observed presence of *nosZ* genes in DNRA-catalyzing microorganisms.

**Importance:** Dissimilatory nitrate/nitrite reduction to ammonium (DNRA) is a microbial energy-conserving process that reduces NO_3_^−^ and/or NO_2_^−^ to NH_4_^+^. Interestingly, many DNRA-catalyzing microorganisms possessing *nrfA* genes harbor *nosZ* genes encoding nitrous oxide reductases, i.e., the only group of enzymes capable of removing the potent greenhouse gas N_2_O. Here, through a series of physiological experiments examining DNRA metabolism in one of such microorganisms, *Bacillus* sp. DNRA2, we have discovered that N_2_O may delay transition to DNRA upon an oxic-to-anoxic transition, unless timely removed by the nitrous oxide reductases. These observations suggest a novel explanation as to why some *nrfA*-possessing microorganisms have retained *nosZ* genes that had probably been acquired via horizontal gene transfers: to remove N_2_O that may otherwise interfere with the transition from O_2_ respiration to DNRA.

## Introduction

Dissimilatory nitrate/nitrite reduction to ammonium (DNRA) is the respiratory reduction of NO_3_^−^ and/or NO_2_^−^ to NH_4_^+^ (1–3). All DNRA-catalyzing isolates examined thus far utilize organic compounds as the source of electrons, although recent culture-independent observations suggest the existence of lithotrophic DNRA in the environment (4, 5). As DNRA and denitrification essentially share the same electron donors and acceptors, the two respiratory NO_3_^−^/NO_2_^−^ pathways compete in the environment, and this competition is often viewed in the context of the relative availability of organic carbon and NO_3_^−^ /NO_2_^−^ (6–10). As DNRA theoretically yields larger amount of energy per molecule of NO_3_^−^ reduced, it has been hypothesized that DNRA would be competitive in reduced environments, often characterized by high C:N ratios (6–10). This redox- or C:N-ratio-controlled competition between denitrification and DNRA was demonstrated in several pure culture studies of organisms harboring both denitrification and DNRA pathways, e.g., *Shewanella loihica* PV-4, as well as in laboratory studies of complex microbial communities (7,11–14). However, environments where DNRA outcompetes denitrification, thus contributing substantially to the fate of NO_3_^−^ are rarely found, apart from highly reduced and/or sulfide-rich marine sediments (5,15). If artificial stimulation of DNRA activity may be possible, either via biostimulation or bioaugmentation approaches, DNRA would have various environmental applications. Outcompeting denitrification with DNRA has been proposed as a means to improve nitrogen management of agricultural soils, as DNRA activation would reduce the amounts of nitrogen lost via denitrification and leaching (2,16,17). In the wastewater sector, DNRA has been suggested as a complement to the annamox process, as DNRA can reverse excessive nitrification and reduce undesired NO_3_^−^ back to NO_2_^−^ and NH_4_^+^ (18,19). Such attractive potential applications warrant further investigation into the DNRA ecophysiology.

Previously, production of NH_4_^+^ from reduction of NO_3_^−^ and NO_2_^−^ has been verified for multiple soil isolates carrying either *nrfA* or *nirB* (3). While NirB has an assimilatory function in many organisms and thus is not exclusive to DNRA, the physiological function of the cytochrome *c*_552_ nitrite reductase encoded by *nrfA* is limited to the respiratory role in DNRA (3,20,21). Further, the NO_2_^−^-to-NH_4_^+^ turnover in the microorganisms possessing *nirB* but no *nrfA* invariably required a fermentable organic substrate as the source of electrons, suggesting that NO_2_^−^ may be used for NADH regeneration, rather than being the terminal electron acceptor for energy conservation (22). For these reasons, the signature functional gene representing the DNRA pathway has long been the *nrfA* gene, and NirB-mediated NO_2_^−^-to-NH_4_^+^ reduction is probably not a respiratory reaction, despite the NH_4_^+^ release observed with *nirB*-possessing microorganisms lacking *nrfA* (23,24).

One of the unresolved conundrums surrounding the *nrfA* gene is its widespread co-presence with the *nosZ* gene, i.e., the gene encoding the nitrous oxide reductase, in bacterial genomes (25–27). Further, several *nrfA-* possessing and DNRA-catalyzing microorganisms with *nosZ*, e.g., *Wolinella succinogenes, Anaeromyxobacter dehalogenans*, and *Bacillus* spp., have been experimentally verified for their ability to reduce N_2_O to N_2_ (25,26,28). NrfA-mediated DNRA release N_2_O, as was demonstrated with the four *nrfA*-possessing soil isolates examined earlier (3). All four strains released 0.4 - 3.0% of reduced NO_3_^−^ as N_2_O, and *Bacillus* strain DNRA2, the only organism possessing *nosZ*, presumably consumed the N_2_O that it produced, as N_2_O accumulation was observed in the presence of C_2_H_2_, but not in its absence (29). Energy conservation via N_2_O reduction was implied in the observed cell growth in the *W. succinogens* (the *nosZ* variant), *A. dehalogenans*, and *B. vireti* cultures fed N_2_O as the sole electron acceptor together with a non-fermentable electron donor (26,30,31). Apparent from these observations, N_2_O-reducing capability would benefit the DNRA-catalyzing organisms by enabling them to utilize the fugitive N_2_O from DNRA, as well as N_2_O released from other organisms in their habitat (26,32,33). Perhaps, as the *nosZ* genes these organisms harbor mostly belong to the clade II, which, in general, tend to exhibit higher affinities to N_2_O, the possession of *nosZ* and the capability to capitalize on sub-micromolar N_2_O may even be crucial for their survival in environmental niches unfavorable for DNRA in competing with denitrifiers (26,32).

One of the four *nrfA*-possessing soil DNRA isolates obtained in the previous study, *Bacillus* sp. DNRA2, carries a *nosZ* genes and is metabolically capable of DNRA and N_2_O reduction (3). This *Bacillus* strain carries neither *nirS* nor *nirK*, and thus lacks the capability to denitrify, but possesses the *napA* gene encoding the periplasmic nitrate reductase and three non-identical copies of *norB* genes encoding nitric oxide reductases. Here, we focused on elucidating the ecophysiological benefits of being able to reduce N_2_O, apart from utilization of N_2_O for energy conservation. A series of physiological experiments were performed with *Bacillus* sp. DNRA2, in which the effect of N_2_O on its DNRA activity were examined in presence and absence of a NosZ inhibitor C_2_H_2_. The results suggested an inhibitory effect of N_2_O on DNRA activity following an oxic-anoxic transition, which was further examined by reverse-transcription qPCR targeting *nrfA* transcripts of cultures exposed to repeated oxic-anoxic spells. This study provides a previously unrecognized evolutionary explanation for possession of *nosZ* by DNRA-catalyzing microorganisms and discusses its implications to the denitrification-vs-DNRA competition.

## Results

### Effect of N_2_O on DNRA in *Bacillus* sp. DNRA2 batch cultures

The batch experiments performed with *Bacillus* sp. DNRA2 with the four different gas amendments (N_2_ only, N_2_O/N_2_, C_2_H_2_/N_2_, or N_2_O/C_2_H_2_/N_2_; see the methods and materials section for details) showed that the transition from aerobic respiration to anaerobic respiration (NO_2_^−^-to-NH_4_^+^ reduction) was delayed by the presence of N_2_O (Fig. 1). The dissolved O_2_ concentration decreased below the detection limit (0.07 mg L^−1^) within 25 hours. The NH_4_^+^ concentration decreased to <5 μM by 20 h in all cultures, presumably due to assimilation, as the cell concentration increased to an OD_600nm_ value of 0.030±0.002 after O_2_ depletion (the OD_600nm_ data are not shown, as no significant growth occurred after O_2_ depletion). In the controls, 0.98±0.02 mM NO_2_^−^ was stoichiometrically reduced to 0.86±0.05 mM NH_4_^+^ within 60 hours of O_2_ depletion (Fig. 1A). In the cultures amended with N_2_O but not C_2_H_2_, N_2_O was completely consumed within 10 h after O_2_ depletion, before any significant consumption of NO_2_^−^ or production of NH_4_^+^ occurred (Fig. 1B). When the experiment was terminated at 73 h, 1.00±0.01 mM NO_2_^−^ was reduced to 0.91±0.03 mM NH_4_^+^, indicating that DNRA was marginally affected by the initial presence of N_2_O. Inclusion of C_2_H_2_ to the headspace resulted in significant delays in NO_2_^−^ consumption and NH_4_^+^ production (*p*<0.05). In the cultures with the headspace initially containing C_2_H_2_ but no N_2_O, N_2_O production began at 29.5 h, and the amount of N_2_O-N eventually reached 3.57±0.29 μmoles N_2_O-N vial^−1^ at 73 h, accounting for 12.0±1.9% of NO_2_^−^ that had been consumed up to this point (Fig. 1C). Reduction of NO_2_^−^ to NH_4_^+^ was significantly delayed under this incubation condition, and only 3.6±0.3 μmoles NH_4_^+^ vial^−1^ was detected at 73 h. In the cultures to which N_2_O was added along with C_2_H_2_, NO_2_^−^-to-NH_4_^+^ reduction was further inhibited (Fig. 1C). The amount of N_2_O increased from 6.6±0.3 to 10.1±0.2 μmoles N_2_O-N vial^−1^ (16.5±2.2% of consumed NO_2_^−^).

**FIG 1.**
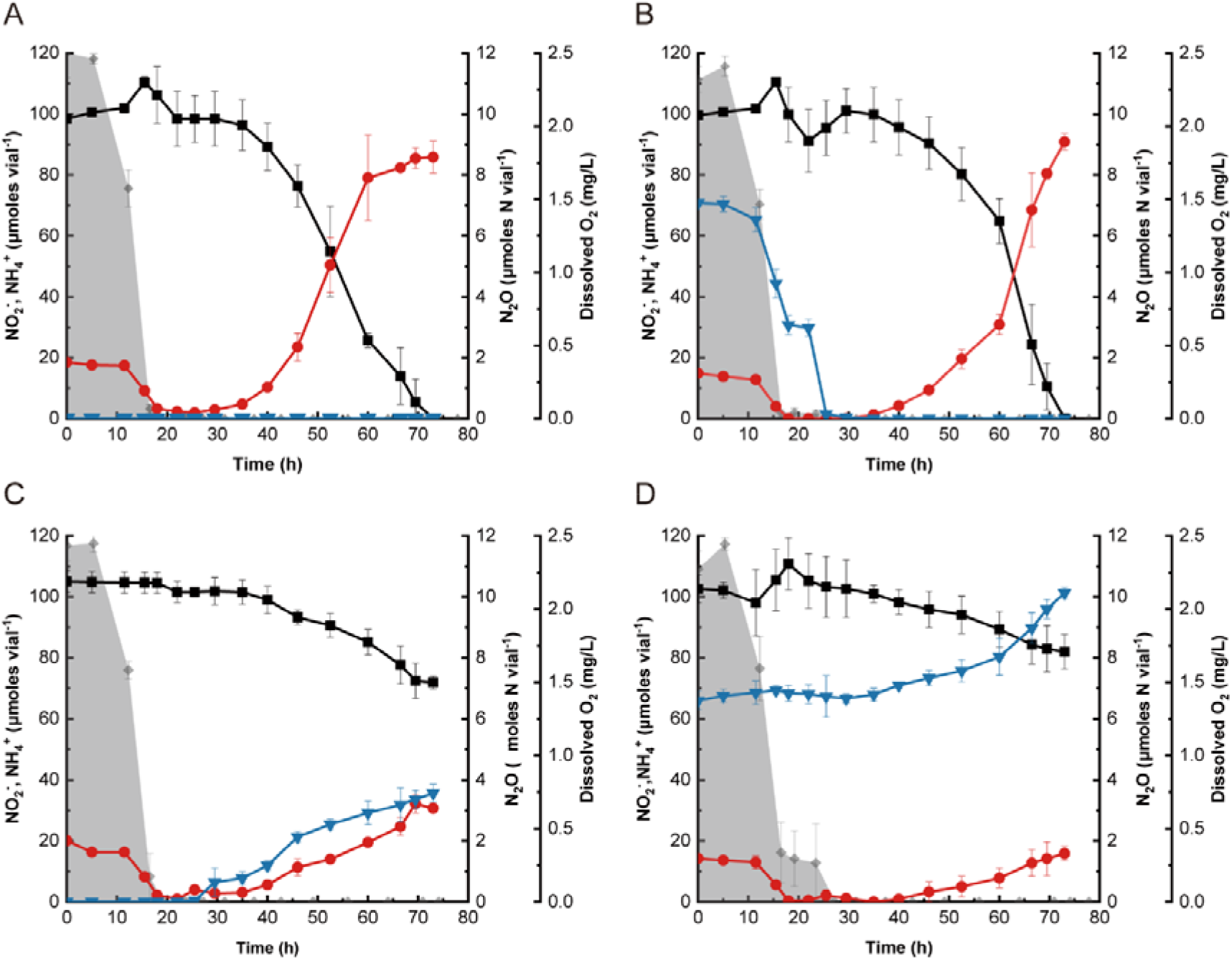
Incubation of 100-mL (prepared in 160-mL serum vials with the headspace consisting of 95% N_2_ and 5% O_2_) *Bacillus* sp. DNRA2 cultures with 1.0 mM NO_2_^−^. The following incubation conditions were examined: (A) control without any headspace amendment, (B) N_2_O-amended condition with 3.5 μmoles N_2_O initially added to the culture vials, (C) C_2_H_2_-amended condition with 10% of the headspace replaced with C_2_H_2_, and (D) N_2_O- and-C_2_H_2_-amended condition. The data points represent the average of triplicate cultures and the error bars the standard deviations of the values obtained from triplicate cultures (■, NO_2_^−^; 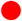, NH_4_^+^; 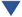, N_2_O-N; shaded curve, dissolved oxygen).

Similar trends were observed when NO_3_^−^ replaced NO_2_^−^ as the electron acceptor (Fig. 2). Without C_2_H_2_ amendment, neither NO_3_^−^-to-NO_2_^−^ nor NO_2_^−^-to-NH_4_^+^ reduction was noticeably affected by the initial presence N_2_O, although N_2_O consumption preceded NO_3_^−^ reduction as the culture turned anoxic (Fig. 2A and B). In the cultures amended with C_2_H_2_, NO_2_^−^-to-NH_4_^+^ reduction that followed NO_3_^−^-to-NO_2_^−^ reduction was substantially slower (Fig. 2C and D). In the cultures amended with C_2_H_2_ but no N_2_O, only 40.4±12.2 of 85.0±11.9 μmoles NO_2_^−^ produced from NO_3_^−^ reduction was further reduced to NH_4_^+^ by the end of incubation, yielding 5.72±0.58 μmoles N_2_O-N. The C_2_H_2_- and N_2_O-amended cultures showed similarly slower NH_4_^+^ production. Only 37.2±0.2 μmoles NH_4_^+^ was produced after 79 h, while the amount of N_2_O increased from 6.1±0.3 to 10.4±0.8 μmoles N_2_O-N vial^−1^.

**FIG 2.**
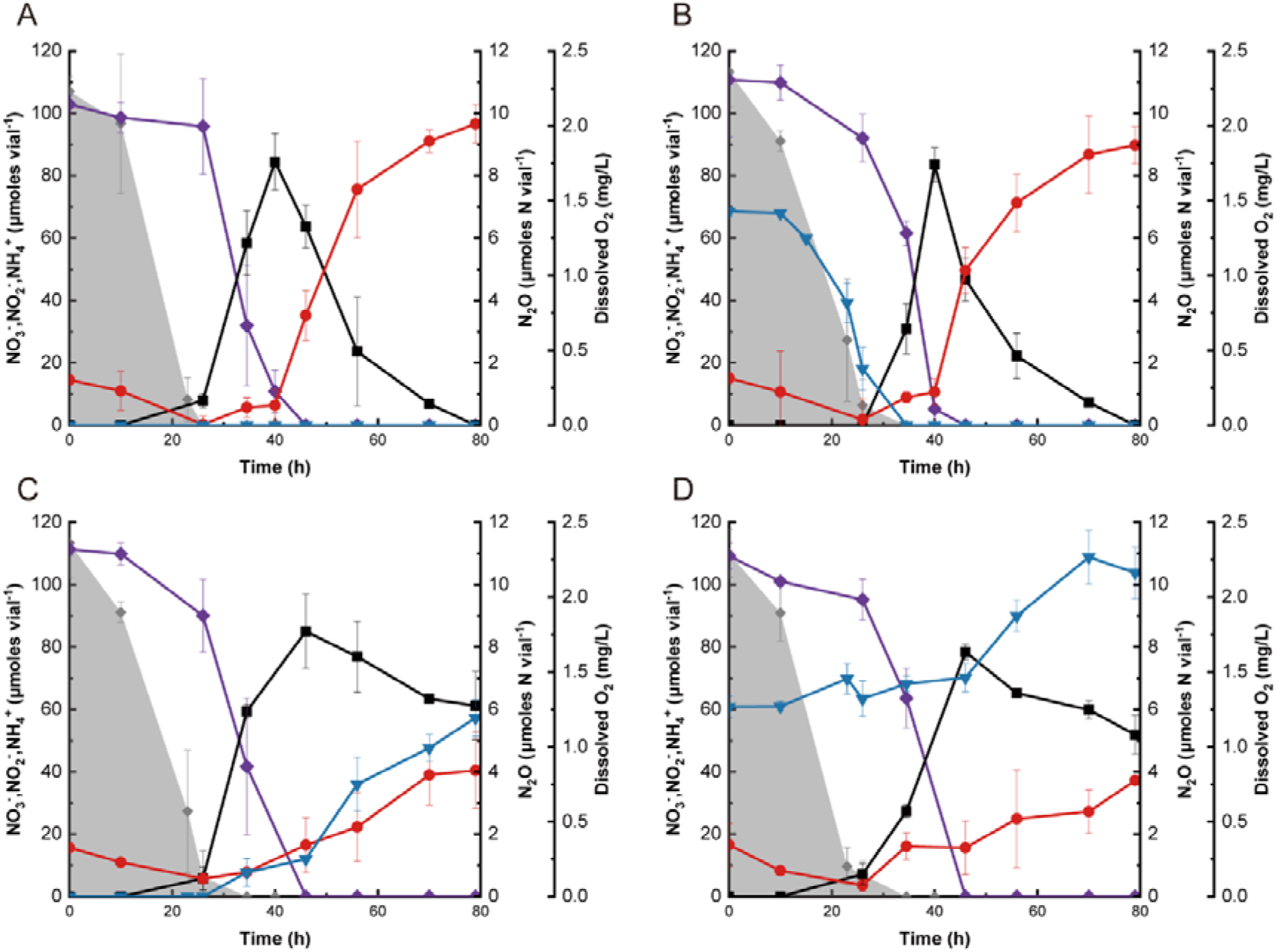
Incubation of 100-mL (prepared in 160-mL serum vials with the headspace consisting of 95% N_2_ and 5% O_2_) *Bacillus* sp. DNRA2 cultures with 1.0 mM NO_3_^−^. The following incubation conditions were examined: (A) control without any headspace amendment, (B) N_2_O-amended condition with 3.5 μmoles N_2_O initially added to the culture vials, (C) C_2_H_2_-amended condition with 10% of the headspace replaced with C_2_H_2_, and (D) N_2_O- and-C_2_H_2_-amended condition. The data points represent the average of triplicate cultures and the error bars the standard deviations of the values obtained from triplicate cultures. (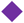, NO_3_^−^;■, NO_2_^−^; 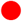, NH_4_^+^; 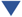, N_2_O-N; shaded curve, dissolved oxygen)

### Confirmation of absence of direct C_2_H_2_ influence on DNRA

The possibility of C_2_H_2_ having contributed to the observed delays in DNRA activation following the oxic-to-anoxic transition was examined in batch reactors fed continuous gas flowthroughs (Fig. 3). In all three reactors, the initial incubation with 3:1 N_2_/air mixed gas increased the cell density to OD_600nm_~0.06. Production of NH_4_^+^ in the reactors began after the gas source was switched to N_2_, N_2_/C_2_H_2_ mixture (9:1), or N_2_/C_2_H_2_ mixture (13:5:2). The NH_4_^+^ production curves of cultures fed with N_2_ were almost identical to those of cultures fed with a N_2_/C_2_H_2_ mixture, while the reactor fed with a N_2_/C_2_H_2_/N_2_O mixture showed substantially slower NO_2_^−^-to-NH_4_^+^ reduction, corroborating the negative impact of N_2_O on DNRA activation. These experiments were repeated with a new set of cultures, reproducing virtually indistinguishable NH_4_^+^ production curves (Fig. S2). These observations substantiated that inhibition of NO_2_^−^-to-NH_4_^+^ reduction observed in the C_2_H_2_-amended batch cultures was most likely due to an inhibitory effect of N_2_O but not of C_2_H_2_.

**FIG 3.**
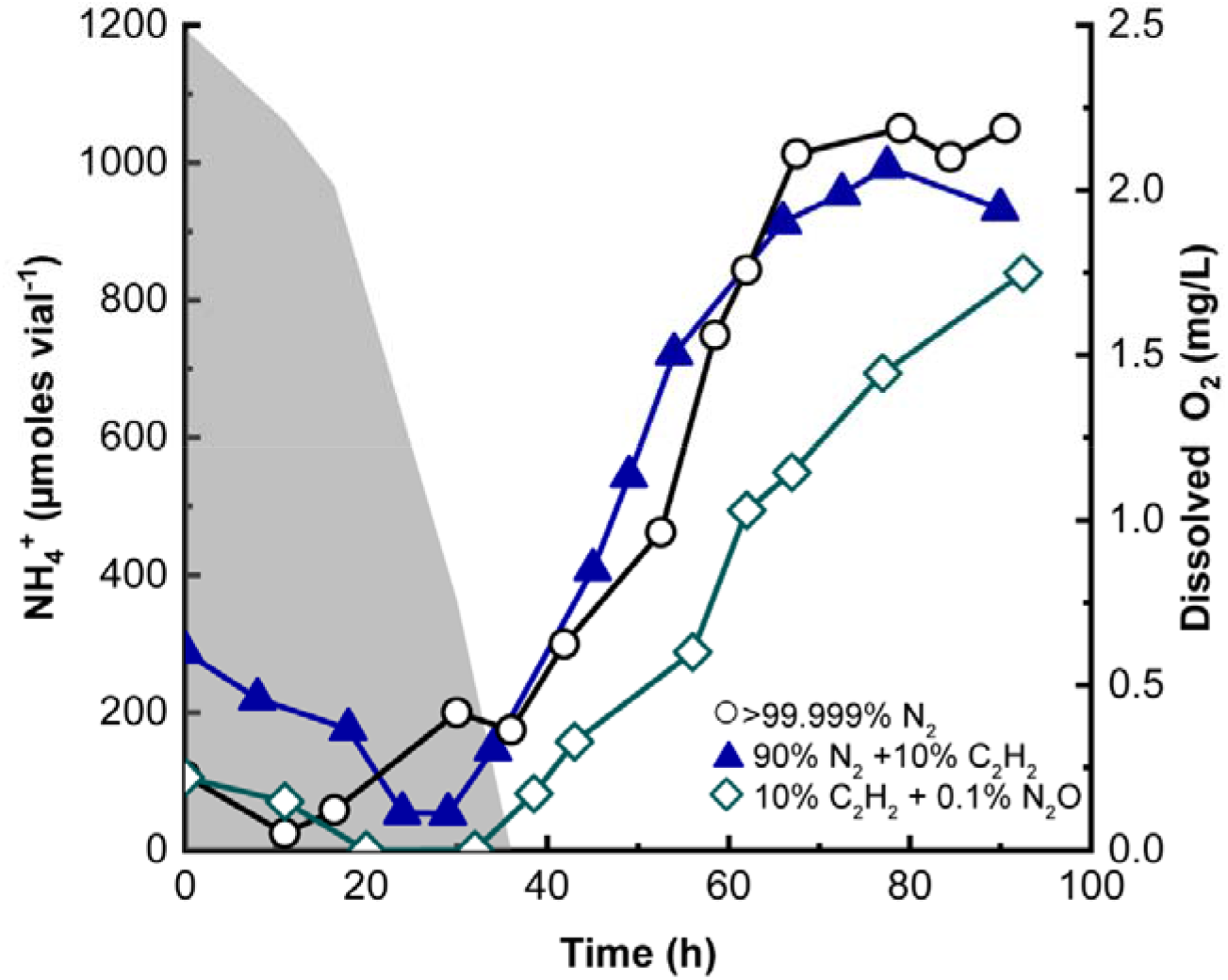
Production of NH_4_^+^ from 2 mM NO_2_^−^ in a batch reactor containing 500-mL *Bacillus* sp. DNRA2 culture fed with continuous stream of >99.999% N_2_ gas (○), 9:1 N_2_/C_2_H_2_ mixed gas (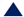), or 9:1 N_2_/C_2_H_2_ mixed gas containing 0.1% N_2_O (◇), after 20 h of aerobic incubation with 95% N_2_ and 5% O_2_. Dissolved oxygen concentration is presented as a shaded curve. The results from the replicate set of experiments are presented in Fig. S2

### N_2_ and NO production during DNRA

Incubation with the closed-circuit robotized incubation system enabled monitoring of NO and N_2_ concentrations in the *Bacillus* sp. DNRA2 cultures amended with and without C_2_H_2_ and N_2_O, showing distinct difference between the two treatments (Fig. 4). The OD_600nm_ values measured after O_2_ depletion were 0.11 ±0.01 and 0.091±0.015 in the controls and C_2_H_2_-and-N_2_O-amended cultures, respectively. The inhibitory effect of C_2_H_2_ and N_2_O on NO_2_^−^-to-NH_4_^+^ reduction was clearly reproducible. The production of 5.8±1.0 μmoles N_2_ (10.7±1.3% of reduced NO_2_^−^) in the controls verified that N_2_O production and consumption occurred simultaneously as NO_2_^−^ was being reduced to NH_4_^+^. The N_2_O yield, in terms of percent of reduced NO_2_^−^, was significantly higher for the cultures amended with C_2_H_2_ and N_2_O, as 17.2±1.8 % of reduced NO_2_^−^ was recovered as N_2_O. Notably, NO accumulated to a substantially higher level in the C_2_H_2_-and-N_2_O-amended cultures than in the controls. While the amount of NO remained below 0.6 nmole vial^−1^ in the controls as NO_2_^−^ was reduced to NH_4_^+^ (44.6-80.6 h), NO accumulated to 2.6±0.8 nmoles vial^−1^ from the background level (~0.5 nmole vial^−1^) in the C_2_H_2_-and-N_2_O-amended cultures between 91.7 and 113.6 h. Evidently, the presence of N_2_O or C_2_H_2_ affected NO production and release in the *Bacillus* strain DNRA2 cultures undergoing DNRA.

**FIG 4.**
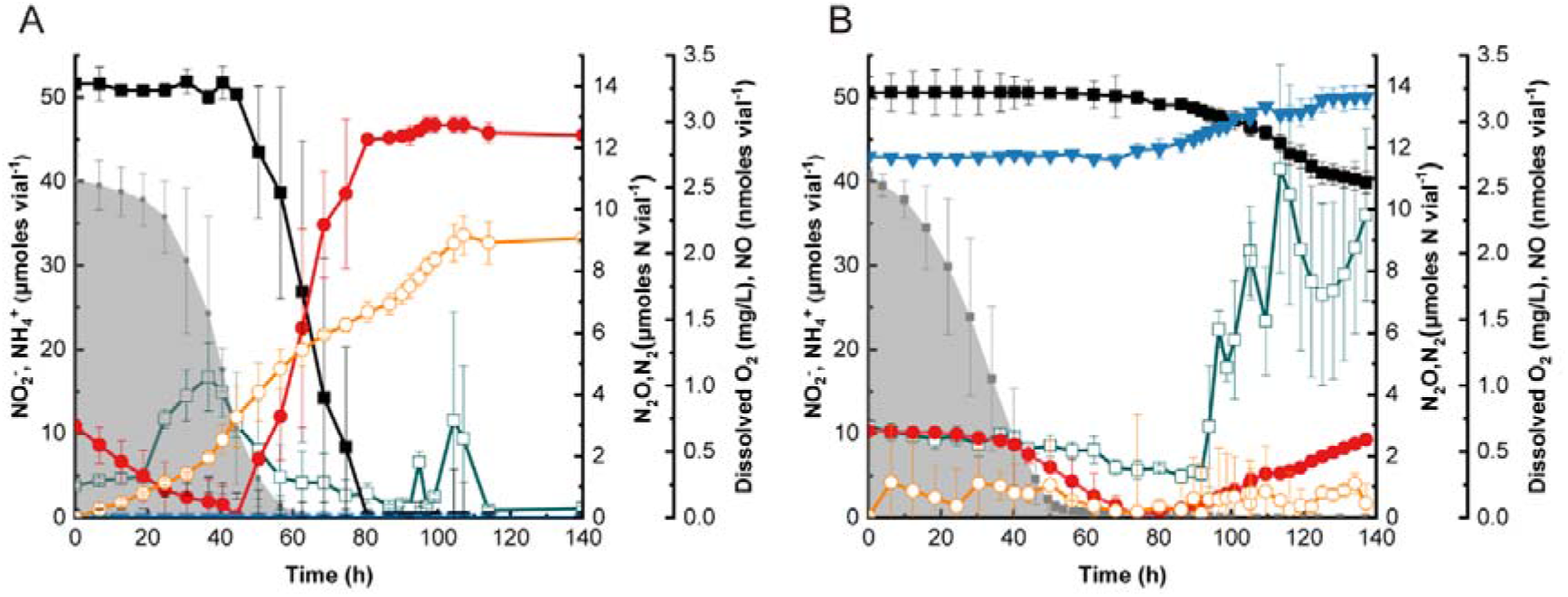
Respiratory kinetics of *Bacillus* sp. DNRA2 batch cultures monitored using a robotized incubation system. Fifty-mL cultures amended with 1.0 mM NO_2_^−^ were incubated in sealed 120-mL glass vials under vigorous stirring. All vials were started with 7 % O_2_ in the headspace. (A) Controls without additional amendments; (B) cultures amended with 6 μmoles N_2_O and 10% (v/v) C_2_H_2_. The data points represent the average of triplicate cultures and the error bars their standard deviations. (■, NO_2_^−^; 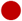, NH_4_^+^; 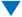, N_2_O-N; 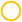, N_2_-N; □, NO; shaded curve, dissolved oxygen)

### Sustained DNRA inhibition in Nos-inhibited *Bacillus* sp. DNRA2 batch cultures subjected to oxic-anoxic alternations

Alternation of oxic and anoxic conditions via periodic replacement of the headspace gas resulted in a more pronounced N_2_O impact on DNRA in *Bacillus* sp. DNRA2 cultures (Fig. 5). In the control, i.e., the batch culture without N_2_O and C_2_H_2_, the vials contained 80.7±5.0 μmoles NH_4_^+^ after 80 h of incubation with 104.0±1.9 μmoles NO_2_^−^ and 12.1±1.2 μmoles NH_4_^+^ as the initial N input. Replacement of the headspace with the oxic gas at 49.5 h immediately halted NO_2_^−^ turnover, which was then recovered after O_2_ depletion. The N_2_O concentration was sustained below the detection limit throughout the incubation, suggesting that produced N_2_O was immediately consumed via NosZ-catalyzed reduction. Each headspace replacement (at 43 and 80 h) resulted in an immediate decrease in NH_4_^+^ concentration, presumably due to assimilation. In the culture vials to which C_2_H_2_ and N_2_O had been added, NO_2_^−^ concentration decreased by only 17.9±3.0 μmoles vial^−1^, while the amount of NH_4_^+^ did not increase above 17 μmoles vial^−1^ at any point during incubation, clearly showing that NO_2_^−^-to-NH_4_^+^ reduction was inhibited to a larger extent with the intermittent headspace replenishment. As the cultures were replenished with a gas containing a high concentration of background N_2_O, no significant increase in the amount of N_2_O could be observed. In all treatments, significant growth occurred only during the initial oxic incubation and immediately following the first headspace replenishment (Fig. S5).

**FIG 5.**
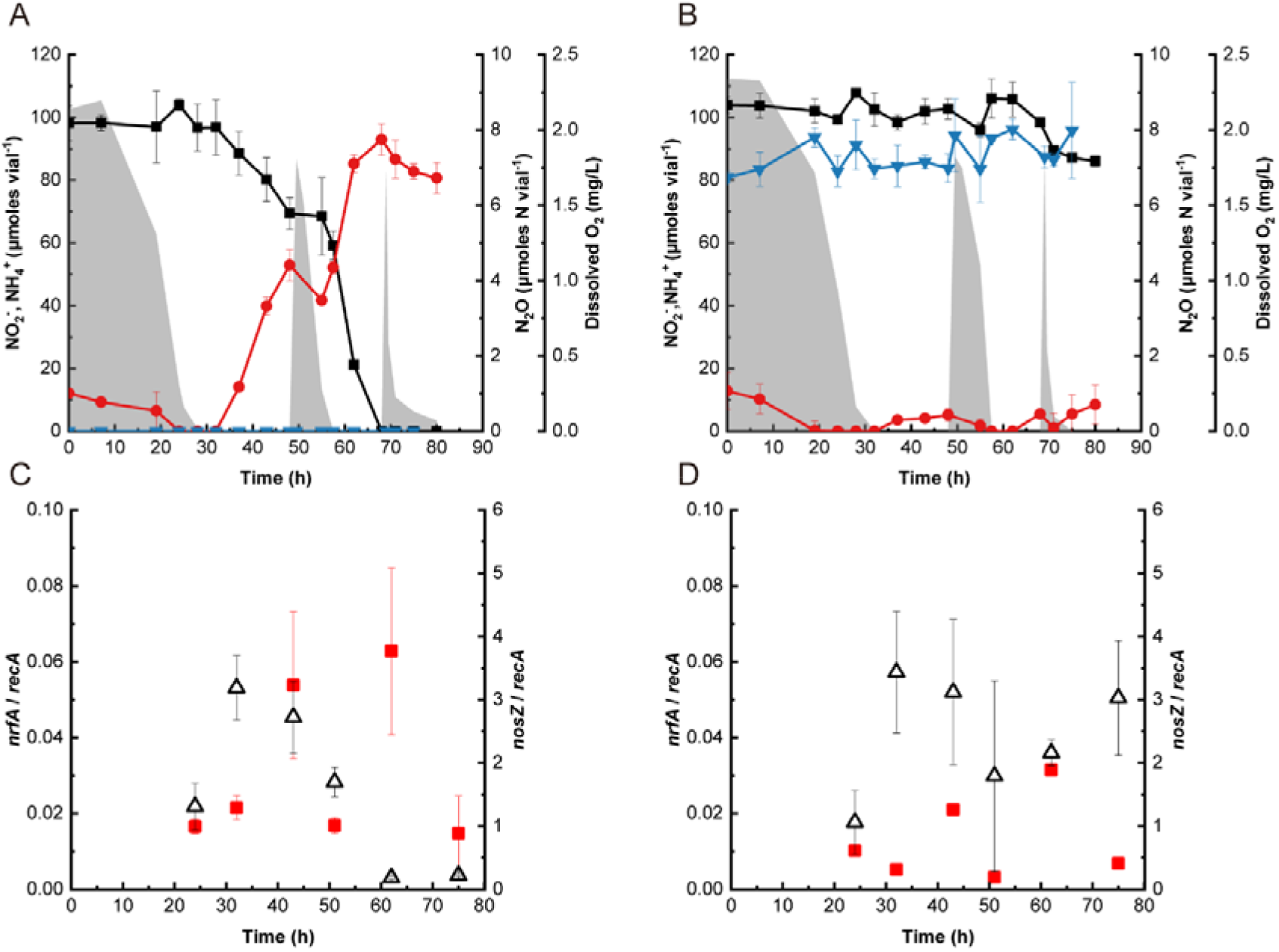
Incubation of *Bacillus* sp. DNRA2 cultures with headspace replenishments to simulate repeated oxic-to-anoxic transitions. The cultures initially contained 1.0 mM NO_3_^−^. The headspace consisted of (A) 3:1 N_2_/air mixed gas or (B) 13:5:2 N_2_/air/C_2_H_2_ mixed gas amended with 3.5 μmoles N_2_O (before equilibration) and was replaced with gas with the same composition at 52.5 h and 70 h. The transcript copy numbers of *nrfA* 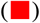 and *nosZ* (Δ) normalized with the copy numbers of *recA* transcripts under the condition A (C) and the condition B (D) were monitored with RT-qPCR. The average values obtained from biological triplicates are presented, and the error bars represent their standard deviations (■, NO_2_^−^; 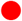, NH_4_^+^; 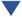, N_2_O-N; shaded curve, dissolved oxygen)

### The effect of N_2_O on *nrfA* and *nosZ* transcription

Reverse transcription quantitative PCR (RT-qPCR) analyses, performed with select samples from the above experiments with repeated oxic-to-anoxic transitions, showed that 15 μM dissolved N_2_O was sufficient to significantly alter *nrfA* transcription in *Bacillus* strain DNRA2 (Fig. 5). With the sole exception of 75 h when NO_2_^−^ had been depleted in the control cultures, *nrfA* transcription was significantly lower for the N_2_O-and-C_2_H_2_-amended cultures than the control cultures (*p*<0.05). The fold-differences between the treatments were substantial, ranging between 1.6 and 5.3. Also notable and common to both sets of cultures, was that the *nrfA* transcription levels measured when O_2_ was present were significantly lower than those measured during the ensuing anoxia (*p*<0.05). For example, the *nrfA* transcription levels measured at 51 h (oxic) for the N_2_O-and-C_2_H_2_-amended cultures and the controls (3.2±2.6 x 10^−3^ and 1.7±0.2 x 10^−2^ *nrfA/recA*, respectively), were both significantly lower (*p*<0.05) than those measured at 62 h (anoxic; 6.3±2.2 x 10^−2^ and 3.1±1.0 x 10^−2^ *nrfA/recA*, respectively). These RT-qPCR results clearly showed that the presence of N_2_O affected *nrfA* transcription in *Bacillus* sp. DNRA2, explaining, at least partially, the inhibition of DNRA activity observed in the N_2_O-and- C_2_H_2_-amended cultures.

The *nosZ* transcription levels were not significantly different between the control cultures and the N_2_O-and- C_2_H_2_-amended cultures until 51 h (*p*>0.05), but were an order of magnitude higher in the N_2_O-and-C_2_H_2_-amended cultures beyond that time point (*p*<0.05; fig. 5b). Apparently, the presence of N_2_O, i.e., the substrate of NosZ, and C_2_H_2_, i.e., a NosZ inhibitor, had no significant effect (*p*>0.05) on *nosZ* transcription. The difference seen towards the end of the experiment may be due to a probable decrease in intracellular NO and/or N_2_O concentration, which would have accompanied NO_2_^−^ depletion. Also notable was that transcription of *nosZ* was at least an order of magnitude higher than that of *nrfA* in both sets, while not being as sensitive to exposure to O_2_ as that of *nrfA*.

## Discussion

Release of N_2_O from NO_2_^−^-to-NH_4_^+^ reduction has been widely observed in DNRA-catalyzing microorganisms, presumably due to leakage of the probable intermediate NO and subsequent reduction by NO reductases (3,25,34–37). While most of these studies have reported N_2_O yields (i.e., mole N_2_O-N released per mole NO_3_^−^ or NO_2_^−^ consumed) below 5%, the yields varied substantially even among phylogenetically close microorganisms (3,35). Further, the experiments with *B. vireti* showed that N_2_O yields may vary depending on growth conditions and also that N_2_O yields may be substantially larger (e.g., >10%) under certain incubation conditions, e.g., high NO_3_^−^ concentration (25,31). For long, DNRA has been perceived as a pathway that yields less N_2_O than denitrification, and the DNRA-catalyzing microorganisms harboring *nosZ* have drawn particular interest as potential net consumers of environmental N_2_O (2,26,27). However, knowing that substantial amounts of N_2_O can be produced from DNRA, the widespread possession of *nosZ* by DNRA-catalyzing microorganisms may need explanations further than that merely pertaining to their N_2_O-scavenging capability.

In *Bacillus* sp. DNRA2, the presence of N_2_O clearly delayed NrfA-mediated NO_2_^−^-to-NH_4_^+^ reduction as the culture was transitioning from aerobic to anaerobic respiration (Fig. 5C and D). The down-regulated *nrfA* transcription in presence of N_2_O explained this delayed onset of DNRA; however, the mechanism via which N_2_O affects *nrfA* transcription remains unelucidated and can only be hypothesized based on the limited observations. Transcription of *nrfA* has been observed only in a surprisingly limited number of DNRA-catalyzing microorganisms, including *Escherichia coli* K-12, *W. succinogens* (the *nosZ_+_* variant), *S. loihica* PV-4, *B. vireti*, and *Citrobacter* sp. DNRA3 (3,7,11,31,38–41). Down-regulation of *nrfA* transcription and DNRA activity in the presence of O_2_ and NO_3_^−^ has been repeatedly observed (3,31,38,41). In *S. loihica* PV-4, harboring both denitrification (with NirK catalyzing NO_2_^−^-to-NO reduction) and DNRA pathways, *nrfA* transcription was significantly affected by the electron donor- or acceptor-limitation, pH, and NO_2_^−^/NO_3_^−^ ratios (7,11,40). The only study that reported N_2_O regulation of *nrfA* transcription was that performed with *W. succinogens nosZ*^+^ variant, where the presence of N_2_O increased transcription of *nrfA*, along with those of *napA* and *nosZ* (39). Nonetheless, none of these fits into the context of the current observation.

Previous studies have repeatedly suggested the role of NO in transcription-level regulations of nitrogen cycling reactions (42–44). NO concentrations above the baseline level were observed in the N_2_O-and-C_2_H_2_-amended culture only after the onset of DNRA; however, the departure from the control culture was evident, in that NO steadily increased with the progression of DNRA (Fig. 3). How N_2_O may alter production and extracellular release of NO or its consumption and detoxification (presumably by NorB and HmpA, respectively) remains enigmatic. Nevertheless, the possibility that NO is involved in N_2_O-mediated down-regulation of *nrfA* transcription should not be neglected, as NO, even at nanomolar concentrations, may act as a signal initiating transcription of denitrification genes (e.g., *nirS, norB*, and *nosZ*) (43, 44, 45,46). A time-series transcriptomic analysis of *Bacillus* sp. DNRA2 cultures during and after transition from aerobic respiration to DNRA would be an interesting follow-up study, in that it may be able to disclose the genes under influence of the N_2_O presence, e.g., transcription regulators, electron transport chain enzymes, and/or even those related to vitamin B_12_ synthesis, that may help elucidate mechanistic features of the N_2_O-elicited gene regulation, including possible NO involvement in the regulatory cascade (47, 48).

In DNRA-catalyzing bacteria possessing *nosZ*, N_2_O reduction is often observed to occur simultaneously with DNRA (3,28). The results from previous experiments with *W. succinogens* (the *nosZ^+^* variants) and *Bacillus* strain DNRA2, which comparatively examined N_2_O evolution in the cultures with and without the N_2_O reduction inhibitor C_2_H_2_, implied that NosZ-mediated N_2_O reduction occurs simultaneously with DNRA in these strains (3,28). Further, in-line N_2_ monitoring verified production of N_2_ from N_2_O reduction during anoxic incubation of *Bacillus vireti* and *Bacillus* sp. DNRA2, as these microorganisms reduced NO_3_^−^/NO_2_^−^ to NH_4_^+^ (25, this study). Utilization of fugitive N_2_O from DNRA as an additional source of electron acceptors would not provide much of an energetic benefit to these organisms. Assuming a 5% N_2_O yield, the additional electron-accepting capacity gained from reduction of the produced N_2_O to N_2_ via NosZ activity would be merely 0.22% of that gained from the dissimilatory reduction of NO_2_^−^ to NH_4_^+^ (See Supplemental Material).

The current study posits a novel hypothesis that *nosZ*-possessing DNRA-catalyzing microorganisms such as *Bacillus* sp. DNRA2 may have retained *nosZ* genes, probably acquired via horizontal gene transfers, to remove N_2_O that would interfere with the transition from O_2_ respiration to DNRA (27). As NO_3_^−^ in the environment is mostly produced from aerobic oxidation of NH_4_^+^, the largest anoxic pools of NO_3_^−^ (and also NO_2_^−^ despite at much lower concentrations) and the most vigorous NO_3_^−^ and NO_2_^−^ reduction activities in soils and sediments are often associated with oxic-anoxic interfaces where O_2_ concentrations fluctuate, and it is likely that micro-niches in such habitats act as hotspots for N_2_O accumulation from nitrification, denitrification, and/or DNRA (49–52). Any substantial delay in the transition to anaerobic respiration would be detrimental for DNRA-catalyzing microorganisms in their competition with denitrifiers (6,7). Hasty generalization should be avoided, as simultaneous occurrence of N_2_O reduction and DNRA has so far been experimentally confirmed only in *Bacillus* sp. DNRA2, *B. vireti*, and *W. succinogens*, and experimental evidence of N_2_O interference with *nrfA* expression has not yet been reported for any microorganism apart from *Bacillus* sp. DNRA2. Examining whether the observed phenotype relating NosZ and NrfA functions can be further generalized would be an interesting follow-up study, which would help further understand evolutionary implication of *nosZ* gene possession by DNRA-catalyzing microorganisms.

*Bacillus* sp. DNRA2, when incubated with N_2_O in the presence of C_2_H_2_ released 12-19% of consumed NO_2_^−^-N as N_2_O-N (Fig. 1d, 2d, and 3b). The only other reported case with >10% conversion of NO_3_^−^/NO_2_^−^ to N_2_O-N in a NrfA-mediated DNRA reaction was that of *B. vireti*, which released up to 49% of reduced NO_3_^−^ as N_2_O-N when amended with 20 mM NO_3_^−^ (25). Under the experimental conditions that resulted in the high N_2_O yields, *Bacillus* sp. DNRA2 and *B. vireti* both showed a lower *nrfA* transcription level and diminished NO_2_^−^-to-NH_4_^+^ reduction rates following an oxic-to-anoxic transition (31). The NrfA enzymes of the two *Bacillus* strains share high level of similarity (73% amino acid level identity). Possibly, the high yields of N_2_O (or of NO, which may have been immediately reduced by nitric oxide reductases or NO detoxification enzymes) may be due to an inherent structural feature of this particular cluster of NrfA. Such high N_2_O yield and N_2_O sensitivity of DNRA may be the rationale for the genomic observations that many of the *Bacillus* spp. harboring a *nrfA* gene in their genomes possess a *nosZ* gene, although verification of this rather bold hypothesis would require further experimental evidences and mechanistic explanations (Table S2) (53).

The N_2_O effects on DNRA, as observed in *Bacillus* sp. DNRA2, may have substantial implications to the fate of nitrogen in the environment. Whether N_2_O-induced delay in *nrfA* transcription and reduced DNRA activity is widely spread among DNRA-catalyzing organisms remains to be investigated, and this phenotype may possibly be limited to *Bacillus* spp. and their close relatives. Even so, *Bacillus* spp. are often an abundant group of microorganisms in agricultural soils, where the fate of NO_3_^−^ has environmental and ecological consequence (54,55). As DNRA-catalyzing microorganisms compete with denitrifiers for the common electron acceptors, i.e., NO_3_^−^ and NO_2_^−^, any delays in NrfA activation or reduced NrfA activity would result in silencing of the DNRA phenotype (3,6,7). In NO_3_^−^-rich microenvironments near oxic-anoxic interfaces in soils, DNRA-catalyzing microorganisms with similar physiology as *Bacillus* sp. DNRA2 would have limited DNRA activities, if local NosZ activity lags behind production or influx of N_2_O. Probably, DNRA-catalyzing *Bacillus* spp. may have retained the *nosZ* genes to increase the chance of competing against denitrifiers in such microenvironments. Supporting that NosZ was playing a crucial role in facilitating DNRA, *Bacillus* sp. DNRA2 was capable of keeping the N_2_O level low and rapidly transitioning from aerobic respiration to NO_2_^−^-to-NH_4_^+^ reduction when incubated without the NosZ inhibitor C_2_H_2_. Going one step further, these NosZ-wielding DNRA-catalyzing microorganisms may be key to collective DNRA enhancement in soils, in that they may provide relief to N_2_O inhibition on DNRA activities of the surrounding microorganisms lacking *nosZ*. Whether and to what extent such hypothetical enhancement to collective nitrogen retention may occur in the soil microbiomes warrant further investigation.

## Methods and Materials

### Culture medium and growth condition

The medium contained per L, 0.58 g NaCl, 0.41 g Na_2_HPO_4_, 0.29 g K_2_HPO_4_, 5.3 mg of NH_4_Cl, 6.2 mg R2A powder (Kisanbio, Seoul, South Korea), and 1 mL 1000x trace metal solution (56). The pH was adjusted to 7.0 with 5 M HCl. Unless otherwise mentioned, batch cultures were prepared with 100 mL medium in 160-mL serum vials. For preparation of anoxic cultures, the vials were flushed with >99.9999% N_2_ gas (Deokyang Co., Ulsan, South Korea) for 15 minutes and sealed with butyl rubber stoppers and aluminum crimps. For preparation of suboxic cultures, a pre-determined volume of the N_2_ headspace was withdrawn and the same volume of air was injected through a 0.2-μm syringe filter (Advantec Inc., Tokyo, Japan). Filter-sterilized 200X vitamin stock was added to the medium after autoclaving (57). Sodium lactate was added to a concentration of 5 mM, and KNO_3_ or NaNO_2_ was added to a concentration of 1 mM unless otherwise mentioned. The medium vials were inoculated with 1 mL of *Bacillus* sp. DNRA2 preculture aerobically grown to the early stationary phase (OD_600nm_ ~0.03). All microbial cultures were incubated in the dark at 25 °C with shaking at 150 rpm, unless otherwise mentioned.

### Batch observation of NO_2_^−^/NO_3_^−^ reduction following oxic-to-anoxic transition

The progressions of DNRA reaction and N_2_O production and consumption were observed in batch cultures of *Bacillus* sp. DNRA2 incubated with NO_2_^−^ or NO_3_^−^ under four different headspace compositions, to examine the possibility that N_2_O may interfere or compete with DNRA reaction (3). Four sets of cultures were prepared: 1) without any amendment to the culturing condition described above; 2) with >99.999% N_2_O gas (Danil Syschem Co., Seoul, South Korea) added to a targeted initial aqueous concentration of 15 μM; 3) with 10% of the N_2_ headspace replaced with >99.99% C_2_H_2_ gas (Special Gas, Inc., Daejeon, South Korea) to inhibit NosZ-mediated N_2_O consumption; and 4) with both N_2_O and C_2_H_2_ added to the aforementioned concentrations. For measurement of the dissolved concentrations of NO_3_^−^, NO_2_^−^, and NH_4_^+^, 1 mL of culture sample was withdrawn, and the supernatant was collected after centrifugation and stored at −20°C. The N_2_O and O_2_ concentrations were measured immediately before the aqueous-phase sampling. The cultures were monitored until NO_3_^−^ and NO_2_^−^ were depleted in the controls (condition 1).

An additional set of batch cultivation experiments was performed to simulate repeated transitions from oxic to anoxic condition and *vice versa* that frequently occur at oxic-anoxic interfaces in soils (51,58). The controls (condition 1) and the cultures amended with both N_2_O and C_2_H_2_ (condition 4) were prepared and the batch incubation experiments were performed identically to the experiments described above but with replacement of the headspace at two different time points during the course of incubation (52 and 70.5 h after start of incubation). Headspace replenishing was performed by flushing the culture vials with N_2_ gas for 5 min and adding, after closure of the culture vials, O_2_, N_2_O, and C_2_H_2_ back to their initial concentrations. The culture samples for reverse transcription (RT)-PCR analyses were collected at 24, 32, 43, 51.5, 62, and 75 h. The *nrfA* and *nosZ* transcripts in *Bacillus sp*. DNRA2 cultures were quantified by RT-qPCR using a previously established protocol (see Supplemental Material for a detailed method) (7).

To isolate the effect of C_2_H_2_ on DNRA from that of N_2_O, NO_2_^−^-to-NH_4_^+^ reduction by *Bacillus* sp. DNRA2 was observed in a fed-batch reactor continuously flushed with N_2_ gas or 9:1 N_2_:C_2_H_2_ mixed gas with or without 0.1% (v/v) N_2_O (Fig.S1). An 1-L glass reactor vessel was prepared containing 490 mL medium amended with 2 mM NaNO_2_, 10 mM lactate, and 0.2 mM NH_4_Cl and inoculated with 10 mL of *Bacillus* sp. DNRA2 culture aerobically grown to OD_600nm_=0.03. The aqueous phase was stirred at 250 rpm using a magnetic bar. Initially, a synthetic gas consisting of ~95% N_2_ and 5% O_2_ was bubbled into the liquid phase of the reactor at 40 mL min^−1^. After 30 hours of incubation, the gas source was switched to N_2_ gas or 9:1 mixture of N_2_ and C_2_H_2_ gas with or without 0.1% N_2_O. Dissolved NO_2_^−^ and NH_4_^+^ concentrations were monitored until no further change was observed.

### Analytical procedures

The gaseous concentration of N_2_O was measured using an HP6890 series gas chromatograph equipped with an HP-PLOT/Q column and an electron capture detector (Agilent, Palo Alto, CA). The injector, oven, and detector temperatures were set to 200°C, 85°C, and 250°C, respectively. The dissolved O_2_ concentration was monitored using a FireStingO_2_ oxygen meter and fiber-optic oxygen sensor spots (Pyroscience GmbH, Aachen, Germany). The total amount of N_2_O in a culture vial was calculated from the headspace concentration using the dimensionless Henry’s constant of N_2_O at 25°C, which was calculated to be 1.68 (59). Dissolved concentrations of NO_2_^−^, NO_3_^−^, and NH_4_^+^ were determined calorimetrically as previously described (60,61). Lactate concentrations were measured using high-performance liquid chromatograph (Shimadzu, Kyoto, Japan) equipped with an Aminex HPX-87H column (Bio-Rad Laboratories, Inc., Hercules, CA) at the start and the end of each incubation to confirm that the initially added amount of lactate was sufficient to deplete all added terminal electron acceptors (data presented in Table S1).

### DNRA and N-oxide production/reduction kinetics measured using a robotized incubation system

*Bacillus* sp. DNRA2 cultures were incubated in a robotized incubation system with frequent monitoring of O_2_ and relevant N-species, with particular interest in NO and N_2_, which were not monitored in the other experiments described in this study. The analyses were performed as previously described with minor modifications (25, 62). Briefly, aerobic pre-cultures, raised under vigorous stirring (600 rpm) using magnetic bars were transferred to sealed 120 ml medical flasks containing 50 mL of the R2A medium described above, to an initial OD_600_ of ~0.03. The medium was supplemented with 0.2 mM NH_4_Cl and 1.0 mM NaNO_2_. Prior to inoculation, the flasks had been made anoxic by repeated He-flushing after which 5 mL O_2_ (7% in the headspace) was added with or without 0.15 mL N_2_O (appr. 12 μmoles N_2_O-N) and 12 mL C_2_H_2_. The cultures were incubated at 25 °C with vigorous stirring. Concentrations of the gaseous compounds were monitored automatically with a TRACE 1310 GC (Thermo Fisher Scientific, Waltham, MA; O_2_, CO_2_, N_2_O, and N_2_) and a NOA 280i Sievers nitric oxide analyzer (Zysense, Weddington, USA) connected to the incubation system.

Aqueous samples for measurements of NO_2_^−^ and NH_4_^+^ concentrations and OD_600nm_ were manually withdrawn. Concentrations of NO_2_^−^ were measured as described previously (25). NH_4_^+^ concentration and OD_600_ were performed as described above.

### Statistical analyses

All experiments, unless otherwise mentioned, were performed in triplicate. Two-way t-tests were used to determine the statistical significance of the pairwise comparisons between two different treatments and one-sample t-test to determine the significance of temporal changes in the transcript copy numbers and the concentrations of the N-species. All statistical tests were performed using R software version 3.5.1 (RStudio Team 2018). The *p*-values lower than the 0.05 threshold were considered significant.

## Acknowledgement

This work was financially supported by the National Research Foundation of Korea (NRF) (Grant No. 2020R1C1C100797013 and 2022R1A4A503144711) and also, in part, by the Research Council of Norway (Project No. 325770).

## Supplemental material

TextS1. Reverse transcription quantitative PCR (RT-qPCR) method

TextS2. Stoichiometric calculation of additional electron-accepting capacity gained by NosZ activity.

TableS1. Lactate concentrations measured before and after incubation

TableS2. Inventories of *nrfA/nosZ* genes in sequenced *Bacillus* genomes

TableS3. Primer sets used for RT-qPCR analyses

FigureS1. Schematic depiction of the batch reactor used for isolating the effect of C_2_H_2_ from that of N_2_O.

FigureS2. Independent replicate of the batch reactor experiment, the result of which was presented in Figure 3.

FigureS3. Microbial growth monitored during the batch incubation experiments performed with headspace replenishments (presented in Figure 3).

